# Age determination through DNA methylation patterns of fingernails and toenails

**DOI:** 10.1101/2021.12.15.472741

**Authors:** Kristina Fokias, Lotte Dierckx, Wim Van de Voorde, Bram Bekaert

## Abstract

Over the past decade, age prediction based on DNA methylation has become a vastly investigated topic; many age prediction models have been developed based on different DNAm markers and using various tissues. However, the potential of using nails to this end has not yet been explored. Their inherent resistance to decay and ease of sampling would offer an advantage in cases where post-mortem degradation poses challenges concerning sample collection and DNA-extraction. In the current study, clippings from both fingernails and toenails were collected from 108 living test subjects (age range: 0 – 96 years). The methylation status of 15 CpGs located in 4 previously established age-related markers (ASPA, EDARADD, PDE4C, ELOVL2) was investigated through pyrosequencing of bisulphite converted DNA. Significant dissimilarities in methylation levels were observed between all four limbs, hence limb-specific age prediction models were developed using ordinary least squares, weighted least squares and quantile regression analysis. When applied to their respective test sets, these models yielded a mean absolute deviation between predicted and chronological age ranging from 6.71 to 8.48 years. In addition, the assay was tested on methylation data derived from 5 nail samples collected from deceased individuals, demonstrating its feasibility for application in post-mortem cases. In conclusion, this study provides the first proof that chronological age can be assessed through DNA methylation patterns in nails.

## Introduction

### Forensic age prediction and DNA methylation

In the forensic field, age estimation contributes to the identification of human remains and may help in the identification of perpetrators by limiting the pool of suspects. Various physiological, chemical and molecular methods with diverse accuracy are available to this end [1]. During the most recent years, age predictions based on DNA methylation (DNAm) patterns has emerged as one of the most accurate methods [2, 3] and has thus become a vastly investigated subject. Elaborate epigenome-wide association studies enabled the identification of the specific genome locations of the CpGs that were seen to change with age. Sets of these CpGs, called epigenetic age estimators or clock CpGs, were coupled with a mathematical algorithm to estimate the chronological age (in units of years) of an individual [4, 5]. Numerous publications have been made on age prediction models using only a few up to several hundreds of clock CpGs. In forensics, where sample material is often highly degraded and only available in limited quantities, those models including only a few clock CpGs are most interesting as less sample material is required for analysis. Over the past decade, many such models have been developed with varying prediction accuracy, of which the most accurate ones allow predictions with an error of only 2-4 years [6, 7, 8, 9, 10]. Mostly blood [2, 6, 8, 10, 11], but also other types of sample material, such as saliva [9], buccal cells [10, 12, 13, 14, 15], semen [16, 17, 18], bones [10, 19] and teeth [6, 20, 21, 22] have been investigated to this end.

### Alternative sample materials

Despite the high accuracy achieved from many previously published DNAm age prediction models in blood and other soft tissues, a major challenge remains when post-mortem decay prevents the use of tissues containing high-quality DNA. In such cases, investigators are directed to opt for alternative samples that are less prone to decay [23]. For instance, bones have been investigated as an alternative sample material to predict age [10, 19]. Based on the regression function of Horvath’s skin & blood clock, including 391 CpGs [24], Lee *et al*. obtained a mean absolute deviation (MAD) of 6.4 years for age prediction in bones [19]. A more concise model based on only 6 CpGs located in 4 genetic markers was developed by Woźniak and colleagues, achieving an error of 3.4 years [10]. In addition, several studies have focused on DNAm age prediction in dental tissues, with promising results [6, 20, 21, 22, 25]. The most accurate age estimation was achieved by Giuliani *et al*., who developed several prediction models based on a total of 42 CpGs located in 3 genes (ELOVL2, PENK, FHL2) [20]. Interestingly, the smallest median absolute deviation between chronological an predicted age of only 1.20 years was obtained by combining DNA methylation data of both dental pulp and cementum, rather than from a single tissue.

### Nails as sample material

Although the results from both preliminary and more elaborate studies are promising, collection of bone and teeth samples involves invasive techniques and DNA extraction is a time-consuming procedure [23, 26, 27]. In addition, environmental conditions under which a body decomposes as well as dental disease and treatment can lead to a reduction of the availability of genomic DNA [26]. An alternative, though much less investigated, sample material are nails, which are already routinely used as a DNA source in identification cases when other samples, such as blood, are not readily available [23, 28, 29]. Human nails are highly keratinised structures located on the dorsal side of the most distal phalanges of the hands and feet. The nail plate is continuously replenished by the nail matrix, which is a small area of highly proliferative tissue located at the proximal base of the nail plate [30]. Nail growth is similar to hair growth, where the proliferating keratinocytes are filled with keratin and the genomic DNA is degraded [31, 32]. Fingernails and toenails continuously grow during one’s lifetime with an average of 3 mm and 1 mm/month respectively, unlike hairs, which grow cyclically. On average, it takes fingernails about 6 months to completely grow out, whereas toenails take 12-18 months to do the same [30]. Nails are chemically robust and somewhat resistant to enzyme attack [30, 31]. This stability is mostly due to the high concentrations of keratin present in the nail plate. Keratin is a fibrous protein that contains high amounts of the amino acid cystine, forming large amounts of disulphide cross-links. Apart from the extensive folding of the proteins in the nail – which is supported by the stable disulphide bonds – Van der Waals interactions, hydrogen bonds and coulombic interactions add up to the chemical and physical resistance to outside influence on nails [30]. These features allow them to remain intact post-mortem for a relatively long period compared to blood and soft tissues, such as fat or muscle. Due to the hydrophobic nature of the composing proteins, they are much less affected by hydrolytic degradation, which in turn slows down microbial degradation [31]. Because DNA degradation is an inherent process in the biogenesis of nails, one would not expect non-root nail samples to yield high quantities of DNA. Yet, it has been suggested that entrapped DNA and disintegrated nuclei remain preserved within the keratin structure of the nail [23, 32], allowing STR-typing. As such, numerous studies have been conducted to evaluate the use of nails recovered from decomposed human remains as an alternative DNA source for STR-analysis. Several factors, such as the post-mortem interval, condition of the cadavers (e.g., putrefied, skeletonized, mummified) and location at which they were found (e.g., outdoors, indoors, submerged in water) were taken into account [28, 29, 33, 34, 35]. Piccinini and colleagues reported full autosomal profiles of nails collected from exhumed remains that had been buried for up to 15 years [33]. Other reports have been made where genetic identification was possible of nails collected from corpses with a post-mortem interval of a few days up to several months, or even years, and irrespective of their discovery location or condition [28, 29, 35]. These results demonstrate that nails make for an at least equally effective sample material for identification as bone [28, 29, 33, 35]. Some laboratories even opt for the use of nails instead of bones for this purpose [28, 29]. Whereas a sizeable amount of research has been done for identification objectives, little is known about the epigenetics, or more specifically the methylation status, of DNA extracted from nails. To date, only two research papers, both from the same research group, have been published on this subject [36, 37]. The aim of these studies was to assess the reliability and specificity of the differentially methylated parental allele method in several human tissues, including fingernails. In addition, one publication of an age prediction model was based on DNAm patterns in hair, where a median absolute deviation of 4.15 years and RMSE of 4.92 years were obtained for a Chinese study population[38]. Considering the resemblance between of nails and hair in terms of biogenesis, and thus micro-structural components, these results are promising for age prediction based on DNAm patterns in nails. However, to the best of our knowledge, nails have never been investigated to this end, despite having proven their value in identification when other samples, such as blood, are not readily available. The scope of the current study was therefore to assess whether fingernail and toenails are suitable biological matrices for age estimation based on DNAm status. To this end, the methylation status of four previously identified multi-tissue age-associated DNA methylation markers (ASPA, EDARADD, ELOVL2 and PDE4C) was investigated and used to develop a model for age estimation based on fingernail and toenail clippings from living individuals. As an indication for its applicability in post-mortem cases, the assay was also tested on a limited set of nail samples from deceased individuals.

## Results

DNA was extracted from nail clippings collected from both hands and feet from 108 living and 5 deceased test subjects. The DNAm status of 15 CpGs located in 4 previously identified age-associated markers (ASPA, EDARADD, PDE4C, ELOVL2) was evaluated through pyrosequencing of the bisulphite converted DNA. The resulting data was subsequently subjected to regression modelling using three distinct methods: ordinary least squares (OLS), weighted least squares (WLS) and quantile regression.

### Correlation with age and differences between sampling locations

DNA was extracted from 433 nail samples, with DNA yield ranging from 2 ng to 2295 ng. 19 samples contained less than 10 ng input DNA, the minimum quantity required to produce accurate DNAm measurements as reported by Naue and colleagues [39], and were therefore excluded from further analysis. DNAm data collection could be completed for 100 samples from the right hand, 99 from the left hand, 95 from the left foot and 98 from the right foot. For each CpG, an assessment was made of whether age, biological gender and sampling location attributed to the variance of methylation measurements. Chronological age was identified as a significant factor in all the investigated loci. Only the methylation status of the CpG located in ASPA showed a significant relationship with biological gender (*p* = 0.007). The effect of location is highly variable between the different markers, as demonstrated in figure 1. Post-hoc analysis revealed that significant differences for multiple CpGs occurred between all four limbs, with the exception of the mutual comparison of both feet. Complete results of the pairwise comparisons between sampling locations are presented in figure S1. The intraclass correlation coefficient (ICC) is a measure to assess the degree of similarity and correlation between two or more datasets. In order to verify whether the methylation levels differed between samples originating from the same subject, 7 ICCs were calculated for each marker at a 95% confidence interval. The interrelation between data from each limb was hereby mutually compared, as well as all four limbs combined (see Figure 2). In general, a higher degree of correlation was discerned in the mutual comparisons of both hands or feet. Conversely, when comparing an upper with a lower limb, the ICCs were appreciably lower. An overview of the exact ICCs, including the 95% interval, is provided in table S1.

**Figure 1.**
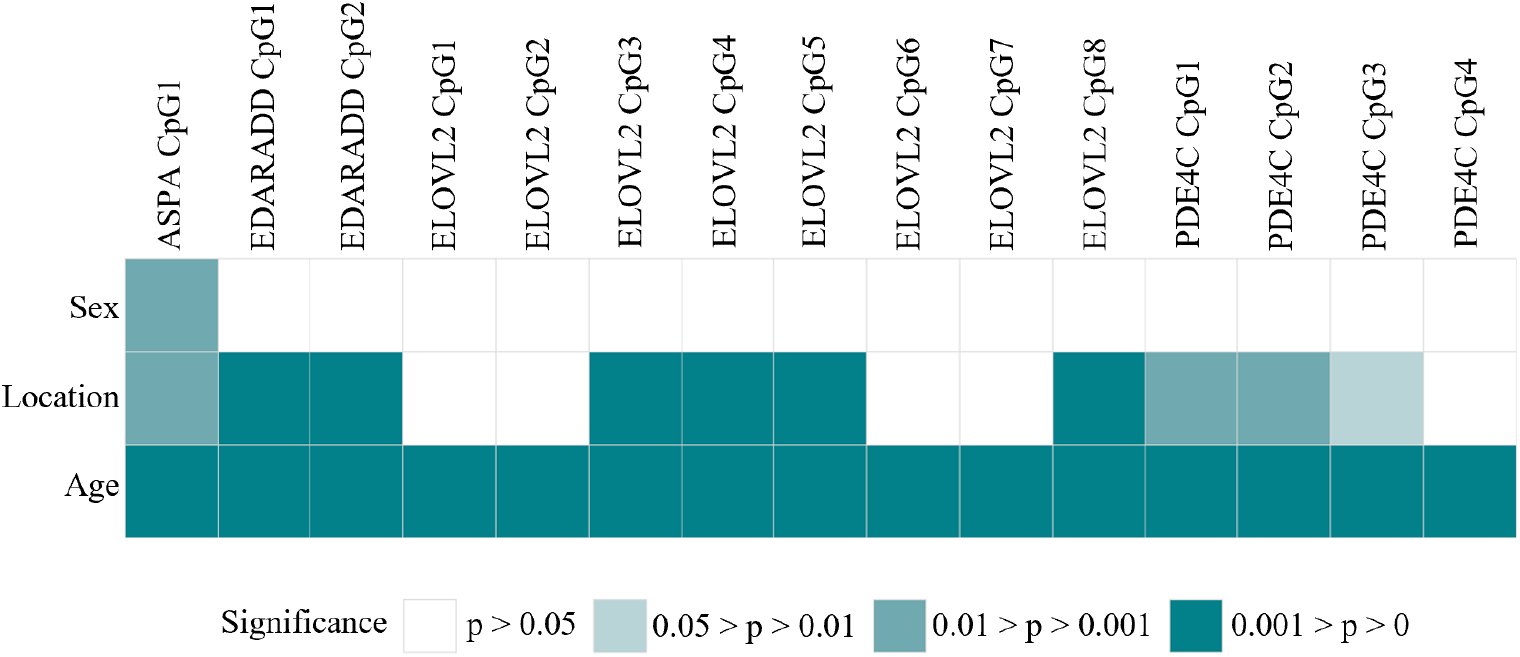
The effects of age, sex and sampling location on the DNAm level of each CpG was tested through a linear mixed-effects regression model.

**Figure 2.**
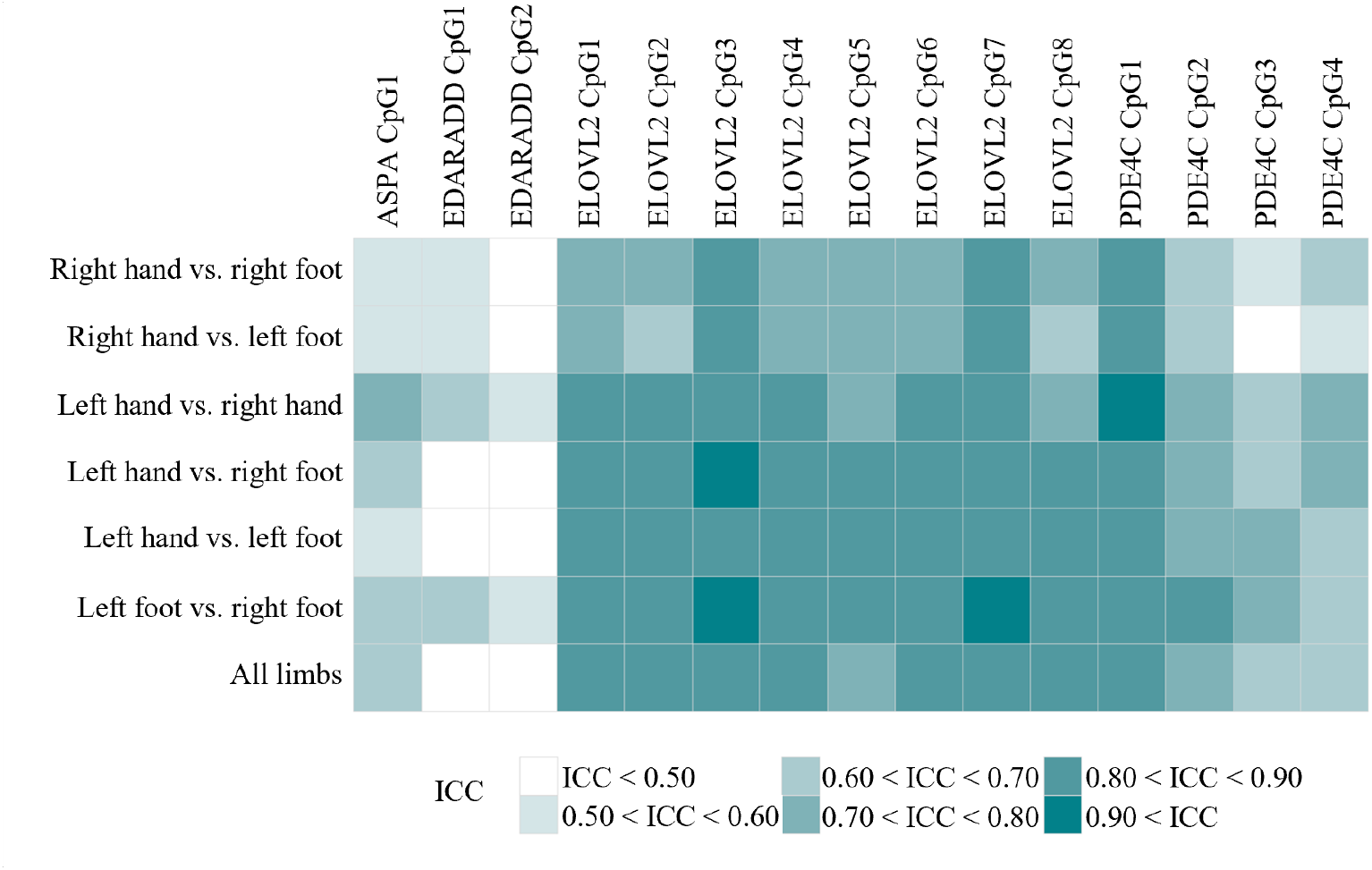
Heat map representing the intraclass correlation coefficients (ICC) calculated for all four limbs combined and for mutual comparison of the four limbs (Two-way mixed effects, absolute agreement, single rater/measurement). A higher ICC indicates a higher degree of correlation and agreement between the compared datasets.

### Age prediction modelling per limb

The correlation between chronological age and methylation levels was evaluated for each limb individually (1). The data was subsequently submitted to regression analysis using three distinctive methods (OLS, WLS and quantile regression). All the investigated CpGs were significantly correlated with age in all four sampling locations (figure 3 and figure S2). Visual inspection of the residual plots and assessment of the 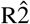 indicated a quadrilinear relationship between age and DNAm measurements for the CpGs located in ELOVL2. The squared values of the methylation measurements were therefore included in the following variable selection. The selected predictors were used to develop an OLS, WLS and quantile regression model for each limb, as shown in **??** and Table S2. Each regression curve is accompanied by a 95% prediction interval, which progressively widens with increasing age in WLS and quantile regression. In order to validate the models, each dataset was split into a training 70% and test 30% set. The regression models were re-fitted on the training sets, which were then applied to the test sets. This resulted in the respective graphs and corresponding parameters presented in figure 4, figure S3 and figure S4. Gender-related DNAm has previously been reported [5, 40] and was also observed for the CpG located in ASPA in the current study. Nevertheless, comparison of the MADs between chronological and predicted age of males and females revealed no significant differences (Wilcoxon rank-sum test *p* > 0.05).

**Figure 3.**
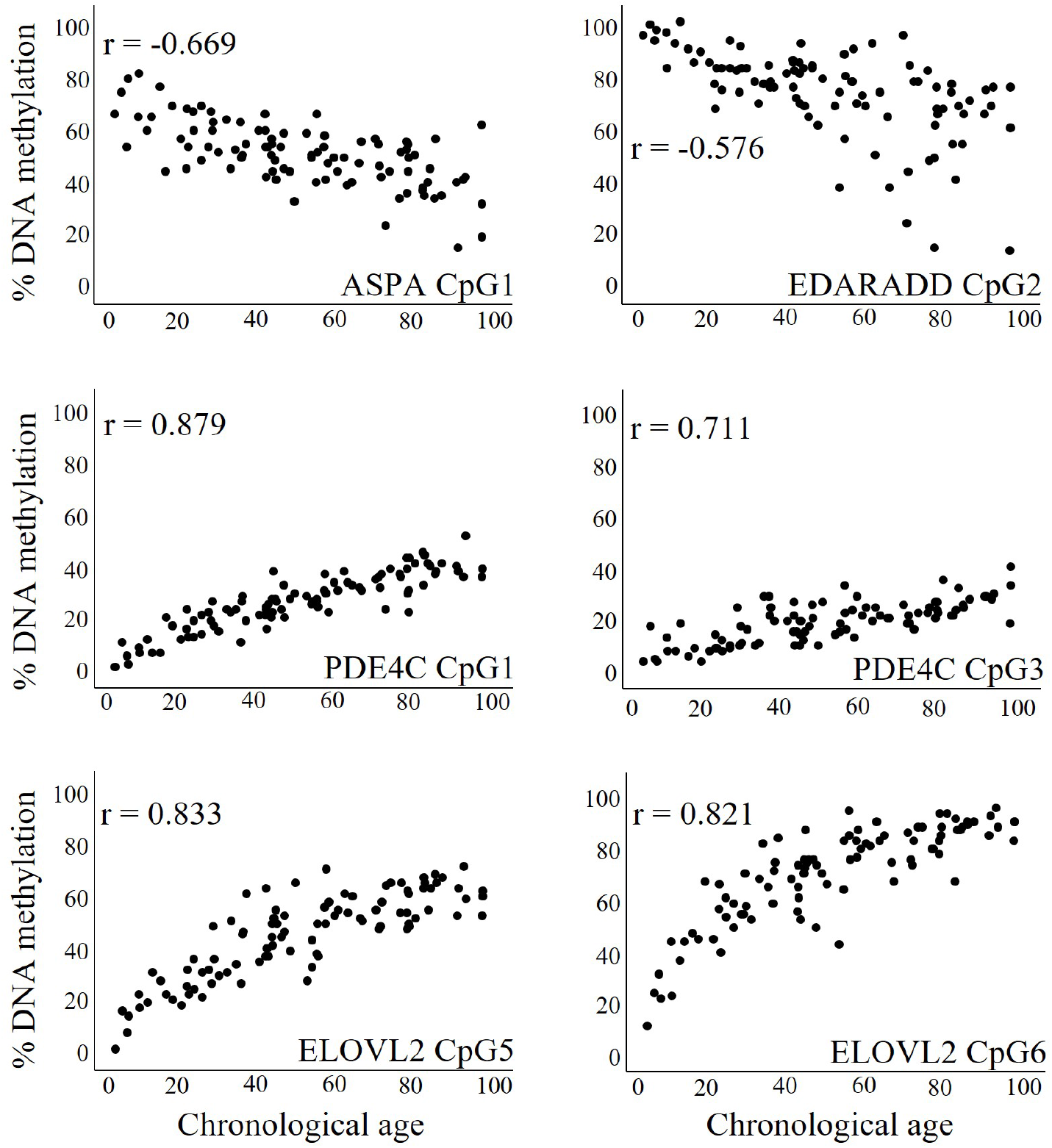
Correlation between DNA methylation and age for the 6 predictors included in the model developed for the left foot dataset.

**Figure 4.**
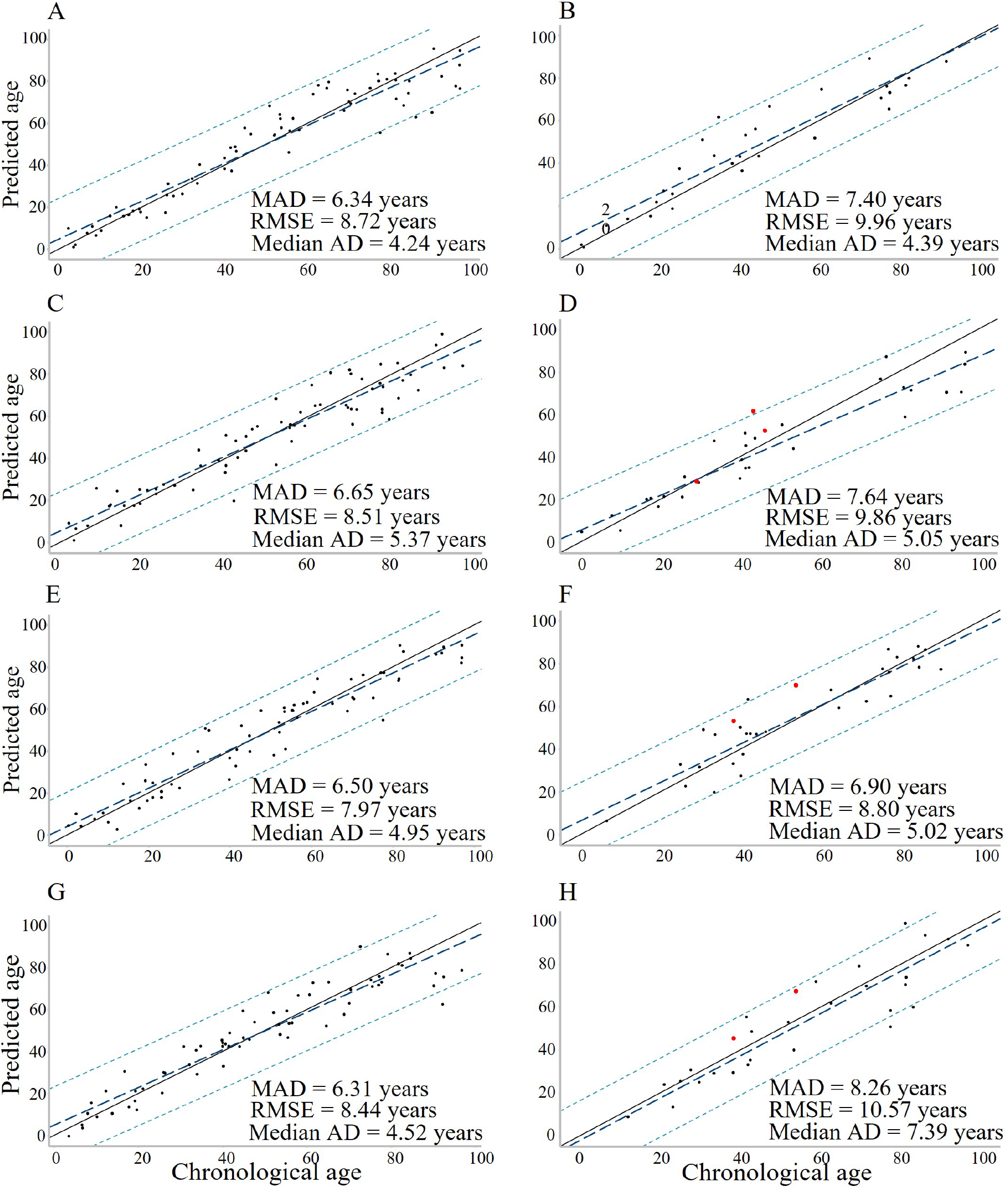
Predicted *vs*. chronological age in the left hand (A, B), right hand (C, D), left foot (E, F) and right foot (G, H) training and test sets respectively. For each sampling location, a different set of prediction variables was selected to develop an OLS prediction model. Red dots indicate predictions made for deceased individuals

DNAm data was obtained from 5 samples from deceased individuals and applied to the prediction model corresponding their sampling location. Because it was not specified whether the toenail samples were taken from the left of right foot, predictions were made using both respective models. All but one prediction fell within the 95% prediction interval in the OLS models (figure 4).

### Limb-specific age prediction models

The current prediction models imply that the sampling location is known, but this is not always the case in practice. Therefore, each training model was tested with the datasets of the other three respective sampling locations to check whether the prediction accuracy would be significantly affected. Significant differences in prediction errors were mostly, yet not exclusively, observed when comparing an upper with a lower limb (figure 5). To investigate whether it would be possible to predict the sampling location based on the observed differences in the DNAm levels, a multinomial logistic regression model was developed. Stepwise regression using the Bayesian information criterion (BIC) resulted in the selection of 5 predictors, allowing to estimate from which limb a sample originated with an overall success rate of 37.91%. Likewise, a model was built to predict whether a sample was taken from either a hand or a foot, regardless of left or right. This model included 3 variables and yielded a success rate of 64.12%.

**Figure 5.**
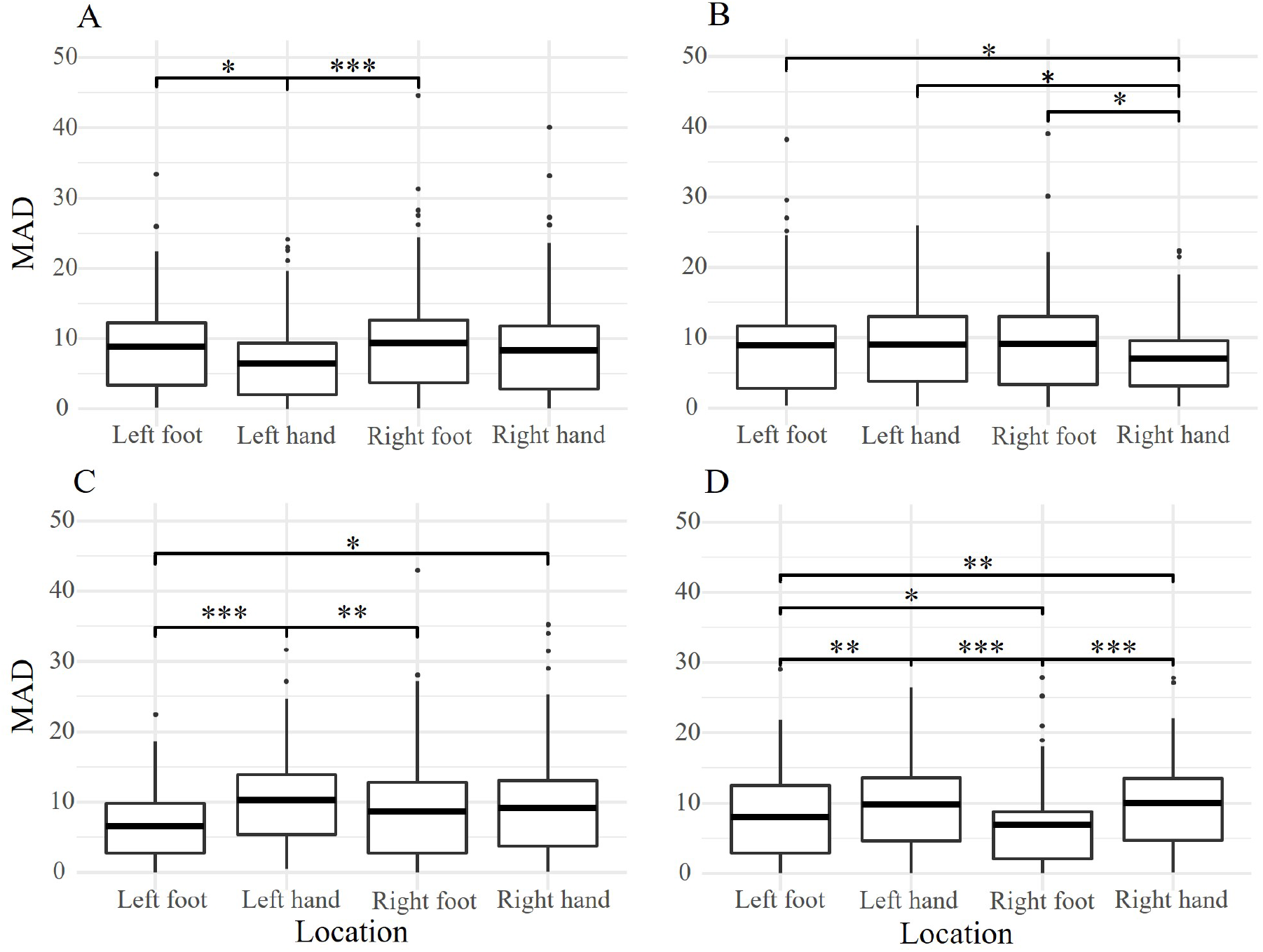
All four datasets were applied to each OLS training model. The resulting absolute prediction errors for the left hand (A), right hand (B), left foot (C) and right foot (D) prediction models were compared through a Wilcoxon signed-rank test. * 0.05 > *p*-value > 0.01, ** 0.01 > *p*-value > 0.001, *** 0.001 > *p*-value > 0

### Comparison with other tissues

Comparison of data obtained from nails (present research) and the data of blood, buccal cells and dentin obtained from teeth [6, 12] revealed that the DNAm status of CpG1 in ASPA, EDARADD and PDE4C and CpG6 in ELOVL2 is tissue-specific. Figure 6 shows the product-moment correlation between age and methylation values of these CpGs for each tissue. The variance of methylation levels could be attributed to both age and tissue type.

**Figure 6.**
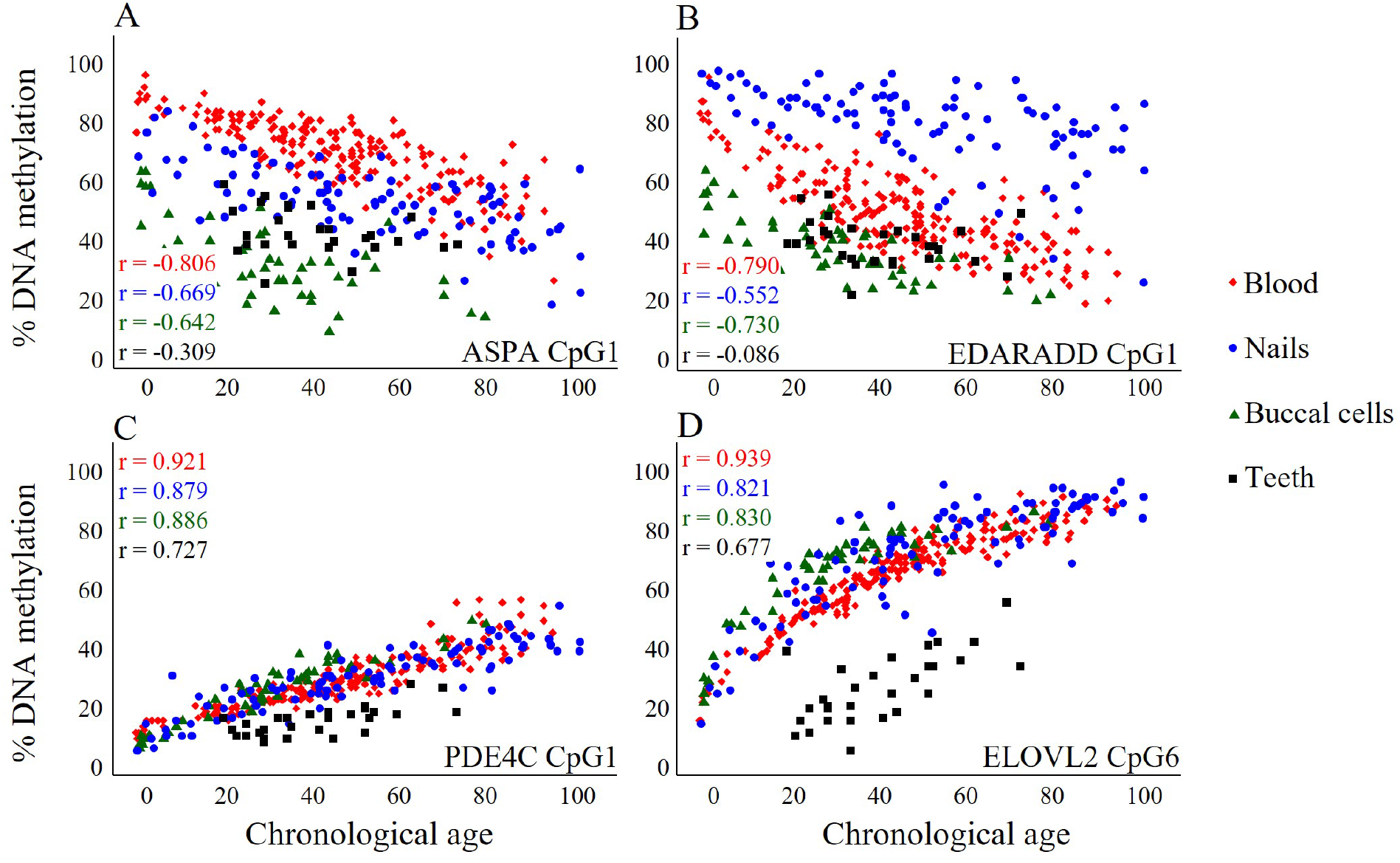
Correlation between DNA methylation and age for CpG1 of ASPA (A), EDARADD (B) and PDE4C (C) and CpG6 of ELOVL2 in blood, nails, buccal cells and dentin extracted from teeth. Note that the data for nails only contains data of the left foot.

## Discussion

To the best of our knowledge, this is the first time that nails have been investigated as a DNA source for the evaluation of the DNAm status and subsequently for age prediction modelling. Data of 15 CpGs located in 4 genes (ASPA, EDARADD, PDE4C and ELOVL2) was collected and used to formulate regression models to predict chronological age.

### Correlation with age and differences between sampling locations

Significant variations were observed in methylation levels between samples originating from different limbs from the same test subject. These deviations were most persistent when comparing hands with feet. To be noted that it takes fingernails about 6 months to completely regrow, where this may take up to 18 months for toenails [30]. This means that the nail clippings provided by the test subjects represented their age subtracted by the time it took for the nails to grow up to the point that they were clipped, rather than their current age. Thus, in theory, this causes a discrepancy of 6-12 months. The most proximal part of a whole nail should be investigated to circumvent this variation, but with the bulk of the test population being healthy volunteers, this was not feasible in the current study. Apart from this source of inconsistency, it is a well-established fact that certain biological and environmental factors also affect epigenetic ageing [7, 41]. A rich body of literature is available on this topic, where the effects of factors such as smoking, BMI and diabetes have been investigated in various tissues [5, 7, 14, 41, 42]. Yet, certain factors could affect the nails on a more local level. For instance, abnormalities of the nail apparatus caused by occupational hazards, such as excess use of irritants or inadequate protection of the hands, have been described [43]. In addition, certain pathogens specifically target nails [32, 43, 44]. For one, onychomycosis has been reported as being the most common nail disease worldwide [45], leading to the question whether it might affect the results of the assay. However, no statistically relevant differences were observed in the comparison of age predictions for affected and healthy nails (Wilcoxon rank-sum test, *p* > 0.05). In future research, additional information of the test subjects, such as hand dominance and occupation, could provide further insights on the occurrence of the observed dissimilarities between the four sampling locations.

### Age prediction modelling per limb

Several studies have recounted an increase of the prediction error with increasing age [6, 14, 25, 38, 46], which was also observed in the present data. OLS regression is the most frequently used statistical model for age prediction, but in case there is evidence of heteroscedasticity of the prediction errors, other methods have been suggested [25, 47, 48]. Predictions were therefore also made using WLS and quantile regression, resulting in similar absolute error parameters as with OLS regression. In contrast, the prediction intervals become larger with increasing age in the WLS and quantile regression models, rather than parallel to the regression curve such as in the OLS model. This allows to report age estimations with a prediction interval appropriately adapted to the occurring variance across different age groups, i.e. younger *versus* older individuals. Note that the prediction interval of the test set of the left foot dataset becomes narrower with increasing age in the quantile regression model. This is most probably coincidental due to the very limited number of randomly appointed test subjects with an age lower than 30 years.

The fact that nails are resilient to degradation and easy to collect would make their use for age prediction most beneficial in post-mortem cases. Therefore, the assay was tested on 5 samples taken from deceased individuals. The results of these preliminary data provide the first indication that it is indeed feasible to use the current methods for its intended purpose of permitting age prediction in human remains. This agrees with other studies, where age estimations have been made based on data from tissues obtained from deceased individuals [6, 10, 15, 19, 46]. Future research with larger sample sizes is required to confirm these initial findings and could provide additional insights on whether the assay would be affected by a longer post-mortem interval and variable post-mortem conditions.

When applying the datasets from each sampling location to the four respective models, the comparison of prediction errors yielded significant differences between all four sampling locations. In practice, this implies that it is necessary to know from which limb a sample was taken in order to provide the most accurate estimations using the correct model. The question was therefore raised whether it would be possible to predict the sampling location based on the DNAm data. With a success rate of only 37.91%, it was not possible to accurately distinguish all four sampling locations from each other. However, based on the DNAm values of only 3 CpGs, it was possible to predict whether a sample originated from either a hand or a foot with a success rate of 64.12%.

By means of comparison, model performances have been surveyed for various tissues that remain (relatively) intact after post-mortem degradation, such as bones [10, 19]), teeth [6, 20, 21, 22, 25] and hair [38]. Where both preliminary and more elaborate data from teeth and bones have yielded MADs ranging roughly from 1.5 to 8 years [6, 10, 19, 20, 21, 22, 25], those obtained from the training and test sets of the current data are rather at the high end of this range. Using hair as sample material, Hao *et al*. reported a median absolute deviation of 4.15 – 6.38 years for their test set, depending on the method of variable selection [38]. These results are comparable with those acquired from the current study.

### Comparison with other tissues

The markers used to produce the models in this study have formerly been investigated in our lab in blood (n = 206), buccal cells (n = 50) and teeth (n = 29) [6, 12]. Not only is there a distinction in the degree of correlation between the various tissues, but there is also dissimilarity between the absolute methylation levels. For instance, for a person aged 25 years, methylation values for CpG 1 in ASPA are ±60% in nails, but only at ±35% in buccal cells. Such variations could most probably be attributed to tissue-specific gene expression [12, 14]. CpG 2 from both PDE4C and ELOVL2 were also included in an assay for age prediction in hair [38]. Despite the previously discussed resemblance between nails and hair, the correlation between DNAm of the locus located in PDE4C and age is strong in the former (r = 0.76 – 0.80), but only moderate in the latter (r = 0.54) tissue. Although such dissimilarities are less distinct in CpG 2 of ELOVL2, the current data revealed a quadratic relationship between DNAm and age, which was not described by Hao and colleagues. These findings might also partly be explained by the fact that their study population only includes a very limited number of subjects aged over 65 years. In contrast, this age group takes up ± 1/3 of the current study population. Another explanation might be found in the previously established influence of ethnicity/ancestry on DNAm patterns [49, 50]. Only one test subject was of Asian origin in the current test population, thus rendering insufficient data to evaluate the effects of ancestry. However, out of the predictions made for the deceased individuals, it was noted to be the only one to fall outside the 95% prediction interval.

### Clinical application

Due to the slow growth rate, non-invasive collection procedure and ease of storage, nails have become a popular matrix in epidemiological studies exploring long term exposure to essential trace metals such as ingested selenium [51] and occupational manganese exposure [52]. Toenails are thereby favoured due to their slower growth rate and lower influence of external contamination compared to fingernails and hair [53]. As many of these types of elements also have been demonstrated to influence the epigenome, clinical studies combining both types of analyses, i.e. trace element exposure and associated DNA methylation changes, seem promising avenues.

## Conclusion

The current study demonstrated for the first time the feasibility of using nails as a DNA source for age prediction modelling. The DNAm status of 15 CpGs located in 4 previously established age-associated genetic markers was evaluated in DNA extracted from nail samples collected from both hands and feet. As significant differences in DNAm levels were observed between all four limbs, limb-specific age prediction models were developed. When each model was applied to their respective test sets, an MAD of 6.5 - 8.5 years was achieved. Although the regression models were developed using samples from living test subjects, testing on a limited number of samples from deceased individuals provided a preliminary indication of their potential use in post-mortem cases. In addition to conventional OLS regression, WLS and quantile regression was conducted to appropriately reflect the increasing prediction error with increasing age. Certain variations, such as variable nail growth, hand dominance and ancestry, were not taken into account in the current study. Future investigation of the effect of such factors and discovery of other age-related loci should allow improvement of the prediction accuracy.

## Materials and Methods

### Sample Collection, DNA extraction and bisulphite conversion

Ethical approval for this study was granted by the Ethics Committee Research UZ / KU Leuven (case number S64520). After obtaining written informed consent, nail clippings from both hands and feet were collected from 108 volunteers (age range: 0-96 years; mean age: 49.42 years; 50-50% women/men) (Fig S4). In addition, 5 samples were collected from deceased individuals, of which 3 from the right hand and 2 from a foot (unspecified whether from left or right foot). The stage of decay of the corpses from which these samples were collected ranged from fresh to advanced decay. All the study subjects were Caucasian, with the exception of one of the deceased individuals, who was of Asian origin. Clippings were pooled per limb and collected in 1.5 ml tubes, which were stored at −20 °C until DNA extraction.

For each individual, all clippings from both hands and feet were used to extract genomic DNA with DNA IQ Casework Pro Kit (#AS1240, Promega) and Tissue and Hair Extraction Kit (#DC6740, Promega) for Maxwell® systems, resulting in 4 DNA extracts per individual. Prior to extraction, the nails were cut with a scalpel into pieces of approximately 5 mm in length after which the samples were incubated in Incubation buffer/Proteinase K at 56 °C for two hours. DNA was subsequently quantified using the Promega PowerQuant® System (#PQ5008, Promega) according to the manufacturer’s instructions.

Extracted DNA (10-200 ng) was bisulphite converted using the Methylamp DNA Modification Kit (#P-1001-2, Epigentek) and eluted in 18 μl elution buffer. Singleplex PCR was then carried out in a total volume of 25 μl, containing 2 μl of converted DNA, 2X Qiagen Multiplex PCR Master Mix (#206143, Qiagen) and 0.8 μM of primers for ASPA, ELOVL2, and EDARADD or 0.4 μM of primers for PDE4C.

### PCR and Pyrosequencing

Table 1 provides an overview of the CpGs that were queried in this study. Note; not all analysed loci have a unique cg number and were therefore all numbered chronologically per marker. Primers and conditions for both DNA amplification and pyrosequencing of four markers (ASPA, EDARADD, PDE4C, ELOVL2) were used as previously described by Bekaert *et al*. [6]. Reproducibility of the pyrosequencing assay was previously demonstrated [6].

**Table 1.**
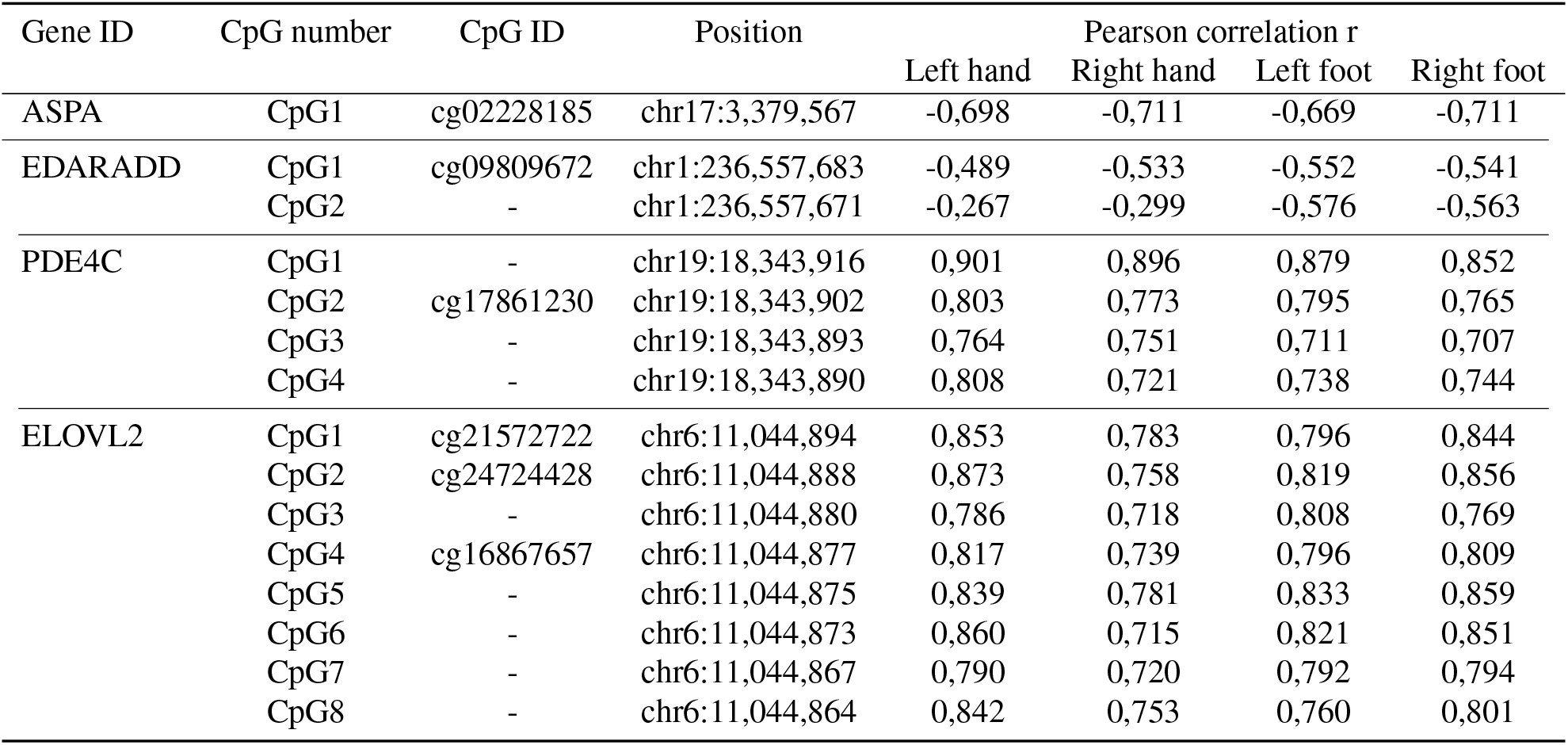
CpGs queried in this study. Genome location and cg number are derived from the GRCh37/hg19 human genome assembly.

| Gene ID | CpG number | CpG ID | Position | Pearson correlation r |  |  |  |
| --- | --- | --- | --- | --- | --- | --- | --- |
|  |  |  |  | Left hand | Right hand | Left foot | Right foot |
| ASPA | CpG1 | cg02228185 | chr17:3,379,567 | -0,698 | -0,711 | -0,669 | -0,711 |
| EDARADD | CpG1 | cg09809672 | chr1:236,557,683 | -0,489 | -0,533 | -0,552 | -0,541 |
|  | CpG2 | - | chr1:236,557,671 | -0,267 | -0,299 | -0,576 | -0,563 |
| PDE4C | CpG1 | - | chr19:18,343,916 | 0,901 | 0,896 | 0,879 | 0,852 |
|  | CpG2 | cg17861230 | chr19:18,343,902 | 0,803 | 0,773 | 0,795 | 0,765 |
|  | CpG3 | - | chr19:18,343,893 | 0,764 | 0,751 | 0,711 | 0,707 |
|  | CpG4 | - | chr19:18,343,890 | 0,808 | 0,721 | 0,738 | 0,744 |
| ELOVL2 | CpG1 | cg21572722 | chr6:11,044,894 | 0,853 | 0,783 | 0,796 | 0,844 |
|  | CpG2 | cg24724428 | chr6:11,044,888 | 0,873 | 0,758 | 0,819 | 0,856 |
|  | CpG3 | - | chr6:11,044,880 | 0,786 | 0,718 | 0,808 | 0,769 |
|  | CpG4 | cg16867657 | chr6:11,044,877 | 0,817 | 0,739 | 0,796 | 0,809 |
|  | CpG5 | - | chr6:11,044,875 | 0,839 | 0,781 | 0,833 | 0,859 |
|  | CpG6 | - | chr6:11,044,873 | 0,860 | 0,715 | 0,821 | 0,851 |
|  | CpG7 | - | chr6:11,044,867 | 0,790 | 0,720 | 0,792 | 0,794 |
|  | CpG8 | - | chr6:11,044,864 | 0,842 | 0,753 | 0,760 | 0,801 |

**Table 2.** Regression formulas of the OLS prediction models developed for each sampling location using their respective training sets. MAD = mean absolute deviation, Median AD = median absolute deviation, RMSE = root-mean-square error.

| Sampling location | Regression model | MAD |  | Median AD |  | RMSE |  |
| --- | --- | --- | --- | --- | --- | --- | --- |
|  |  | Train | Test | Train | Test | Train | Test |
| Left hand | Age (years) = 34.539 - 0.349 (EDARADD CpG1) + 0.988 (PDE4C CpG1) + 0.003 (ELOVL2 CpG2)<sup>2</sup> - 0.657 (ELOV2 CpG6) + 0.009 (ELOV2 CpG6)<sup>2</sup> | 6.34 | 7.40 | 4.24 | 4.39 | 8.75 | 9.96 |
| Right hand | Age (years) = 33.909 - 0.472 (EDARADD CpG1) + 1.404 (PDE4C CpG1) + 0.495 (ELOVL2 CpG2) - 0.425 (ELOVL2 CpG4) + 0.006 (ELOVL2 CpG8)<sup>2</sup> | 6.65 | 7.64 | 5.37 | 5.05 | 8.51 | 9.86 |
| Left foot | Age (years) = 58.832 - 0.247 (ASPA CpG1) - 0.367 (EDARADD CpG2) + 0.896 (PDE4C CpG1) + 0.499 (PDE4C CpG3) + 0.003 (ELOVL2 CpG5)<sup>2</sup> - 0.797 (ELOVL2 CpG6) + 0.008 (ELOVL2 CpG6)<sup>2</sup> | 6.50 | 6.90 | 4.95 | 5.02 | 7.97 | 8.80 |
| Right foot | Age (years) = 30.933 - 0.466 (EDARADD CpG1) + 0.608 (PDE4C CpG1) + 0.575 (PDE4C CpG4) + 0.006 (ELOVL2 CpG2)<sup>2</sup> + 0.002 (ELOVL2 CpG7)<sup>2</sup> | 6.31 | 8.26 | 4.52 | 7.39 | 8.44 | 10.57 |

DNA was amplified under the following conditions: an initial hold at 95 °C for 15 min followed by 40 cycles of 30 s at 95 °C, 30 s at 56 °C, and 30 s at 72 °C for ASPA and EDARADD; 50 cycles of 35 s at 95 °C, 35 s at 52.9 °C, and 35 s at 72 °C for PDE4C and 40 cycles of 30 s at 95 °C, 30 s at 60 °C, and 30 s at 72 °C for ELOVL2. All amplifications ended with a final extension step at 72 °C for 6 min.

In order to perform pyrosequencing, Streptavidin Sepharose High Performance beads (#17-5113-01, GE Healthcare) were used to immobilize 10 μl (ELOVL2) or 20 μl (ASPA, EDARADD, PDE4C) of the biotinylated PCR amplicons. Next, an annealing step with 25 μl of 0.3 μM sequencing primer was performed at 80 °C for 2 minutes, followed by a cooling down period of 10 minutes. Pyrosequencing was performed on the Pyromark Q24 instrument (#9001514, Qiagen), using Pyro Gold Reagents (#970802, Qiagen) according to manufacturer’s instructions. The PyroMark Q24 software v2.0.8 was used to analyse the resulting pyrograms

### Statistical analysis

For each locus, the effects of chronological age, sampling location (left hand, right hand, left foot, right foot) and gender on DNAm levels were determined in a linear mixed-effects regression model. The complete dataset was then split per sampling location and the intraclass correlation coefficient (ICC) was computed for each CpG to assess the degree of similarity and correlation of DNAm levels between all four limbs. Within each dataset, the Pearson correlation between the DNAm status of the CpGs and age was determined.

To account for the observed quadrilinear relationships with age, the squared values of the DNAm levels of the CpGs located in ELOVL2 were included in the collection of variables. A stepwise selection using the Bayesian Information Criterion (BIC) was conducted to select the most suitable combination of predictors for each sampling location. The selected set of variables was used to fit prediction models through ordinary least squares (OLS), weighted least squares (WLS) and quantile regression models. The models were validated by randomly splitting each dataset in a training (70%) and test (30%) set. The selected prediction variables were used to compute an independent regression for the training set, which was validated with the test set. In addition, each model was trained with the datasets of the other three respective sampling locations to test whether there would be a significant difference in prediction accuracy. The resulting MADs were compared using paired Wilcoxon signed-rank tests.

The methylation levels of CpG1 of ASPA, EDARADD, PDE4C and CpG6 of ELOVL2 of the current datasets were compared with those previously acquired for other tissues [6, 12] through analysis of a linear mixed-effects model. All data analyses were performed in R v3.6.1 [54] with RStudio v1.2.5001 [55].

## Acknowledgements

We would like to thank the laboratory technicians at the Laboratory for Forensic Genetics as well as the post-mortem assistants at UZ Leuven for their assistance during the collection and processing of the samples.

## Author contributions statement

Kristina Fokias: Conceptualization, Sample Collection, Methodology, Writing

Ronny Decorte: Commenting on Manuscript

Wim Van de Voorde: Sample Collection, Commenting on Manuscript

Bram Bekaert: Conceptualization, Sample Collection, Methodology, Writing

## Ethics statement

Ethical approval for this study was granted by the Ethics Committee Research UZ / KU Leuven (case number S64520).

## Additional information

The authors declare no conflict of interest.

